# GBAT: a gene-based association method for robust *trans*-gene regulation detection

**DOI:** 10.1101/395970

**Authors:** Xuanyao Liu, Joel A Mefford, Andrew Dahl, Meena Subramaniam, Alexis Battle, Alkes L Price, Noah Zaitlen

## Abstract

Identification of *trans*-eQTLs has been limited by a heavy multiple testing burden, read-mapping biases, and hidden confounders. To address these issues, we developed GBAT, a powerful gene-based method that allows robust detection of *trans* gene regulation. Using simulated and real data, we show that GBAT drastically increases detection of *trans*-gene regulation over standard *trans*-eQTL analyses.

## Main

Identification of long range *trans*-gene regulation often leads to the discovery of important disease-causing genes and pathways that are not captured in *cis* analyses^1-4^. This is because *trans* regulation harbors more cell type specific effects^1, 5, 6^ and explains more than twice the variability in gene expression than *cis* effects^5-7^. However, robust discovery of *trans-*eQTLs is challenging and prone to false positives for four reasons. First, genome-wide scans for *trans*-eQTLs suffer from heavy burden of multiple testing^6, 8^. Second, *trans* effects are typically much smaller than *cis* effects^8^. Third, sequence read mapping errors, multi-mapped reads, and reads from repeat regions, lead to many false *trans* signals^9^. Fourth, naïve use of dimensionality reduction techniques to estimate confounding effects (such as PEER^10^ or SVA^11^) in *trans*-eQTL studies can both reduce power^12, 13^ and introduce false positives in *trans-*eQTL studies^11, 14^.

We address these issues through a new gene-based method, GBAT, for detecting *trans-* regulatory effects. GBAT consists of three main steps (Figure 1). First, to reduce the number of false positives due to mapping issues, GBAT filters out reads that are multi-mapped^15^. In addition, GBAT further removes problematic mapped reads that are not marked as “multi-mapped” by RNA-seq alignment algorithms by discarding reads that are mapped to repeat regions (genomic regions with mappability scores lower than 1, see Supplementary Notes).

**Figure 1.**
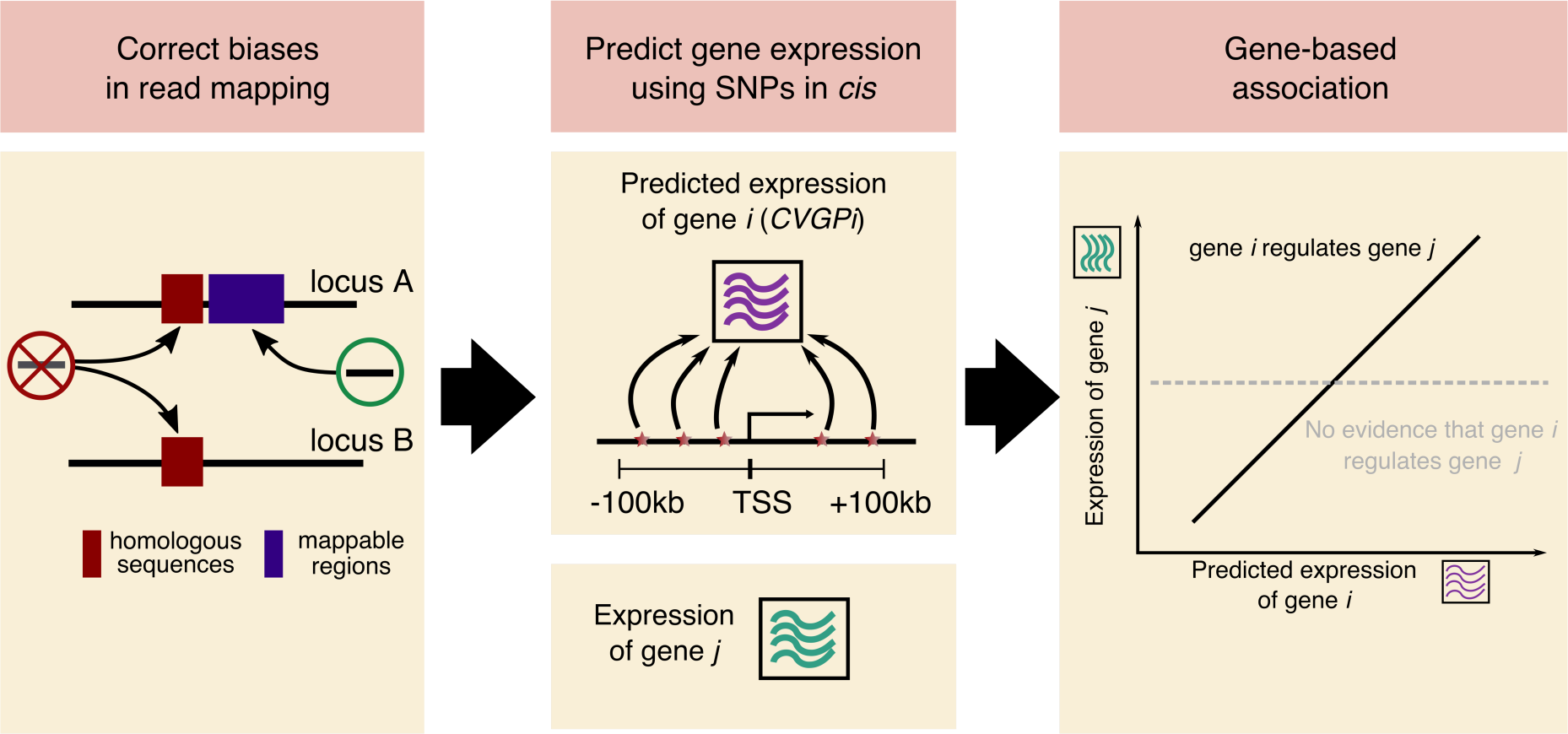
Schematic of the GBAT pipeline. First, GBAT discards reads that are mapped to homologous sequences in the genome in order to prevent false positives in *trans* association mapping. Discarded reads include multi-mapped reads and reads that are mapped to regions of low mappability. After expression quantification, GBAT predicts the the expression levels from *cis* genetic variants (cross-validated cis-genetic prediction of gene i (*CVGP*_*i*_)) using cvBLUP. Finally, GBAT performs gene-based association tests between *CVGP*_*i*_ and expression level of gene *j* to identify gene-based *trans* association, while properly including supervised surrogate variables conditional on each *CVGP*_*i*_ (SV_i_) as covariates.

Second, GBAT uses cvBLUP, a novel gene-based method to produce predictions of gene expression from SNPs *cis* to each gene. The cvBLUP method does not rely on external eQTL studies, but builds leave-one-sample-out cross-validated *cis*-genetic predictions (*CVGP*_*i*_ for each gene *i*), to avoid overfitting issues of standard best linear unbiased predictor (BLUP)(see Methods for details). The cvBLUP method dramatically reduces computing time, compared to other leave-one-sample-out cross validation approaches implemented for prediction methods such as BSLMM^16^ and Elastic-net^17^. This gain is attained by building our *N* (*N*=sample size) leave-one-out *CVGP* predictions after fitting the model only once instead of *N* times (Methods).

Finally, we test the association of each *CVGP*_*i*_ with quantile normalized expression levels *E*_*j*_ of every *trans* gene *j* (at least 1Mb away from gene i). *Cis-*eQTL studies typically include covariates such as PEER factors or surrogate variables from SVA that are intended to model confounders^17, 18^. To prevent false positives^13^ and power loss in *trans-*eQTL studies^11, 12^, the use of supervised versions of PEER and SVA is recommended^13^. Therefore, for each *CVGP*_*i*_, we run supervised SVA conditional on *CVGP*_*i*_, such that the resulting surrogate variables *SV*_*i*_ does not include the genetic effects of gene *i*. We then use *SV*_*i*_ as covariates. While including conditional SVs as covariate is computationally efficient in our gene-based approach, it is impossible in SNP based *trans* association testing.

In addition, GBAT regresses *E*_*j*_ on *SV*_*i*_ and uses the quantile normalized residuals *E*_*j*_^*’*^ for gene-based testing using the regression: *E*_*j*_^*’*^∼ *CVGP*_*i*_. We found that the p-values from this test are well calibrated (Supplementary Notes, Figure S2). However, skipping the normalization caused false positive inflation, as did using the models: *E*_*j*_ ∼ *CVGP*_*i*_ + *SV*_*i*_ or *E*_*j*_ ∼ *CVGP*_*i*_ + *SV* (where *SV* is the naïve SVA not conditioning on *CVGP*_*i*_) (Figure S2).

Using real genotypes from a whole blood RNA-seq dataset: the Depression Genes and Networks cohort^9^ (DGN, sample size N=913), we performed simulations to assess the power of our gene-based approach (GBAT) in comparison to a SNP-based approach (Methods). We simulated a causal SNP→cis-expression→trans-expression model with realistic effect-sizes under different genetic architectures (proportion of causal SNPs = 0.1%, 1% and 10%) and sample sizes (N=200,400,600 and 913) (Supplementary Notes). To better reflect the imperfect genotyping of the individuals, we assumed that only 10% of the SNPs are genotyped, such that not all causal SNPs are observed. We measured the power of both approaches to identify significant *trans* associations (accounting for 1 million×15000 SNP-gene pairs for SNP-based approach or 5,000 ×15,000 (highly heritable gene)-gene pairs for the GBAT approach, using Bonferroni correction). Across all simulated genetic architectures of gene expression and combinations of *cis-* and *trans-* effects, we observed that the power of the GBAT approach is substantially higher than the SNP-based method (Figure 2a, Figure S3).

**Figure 2.**
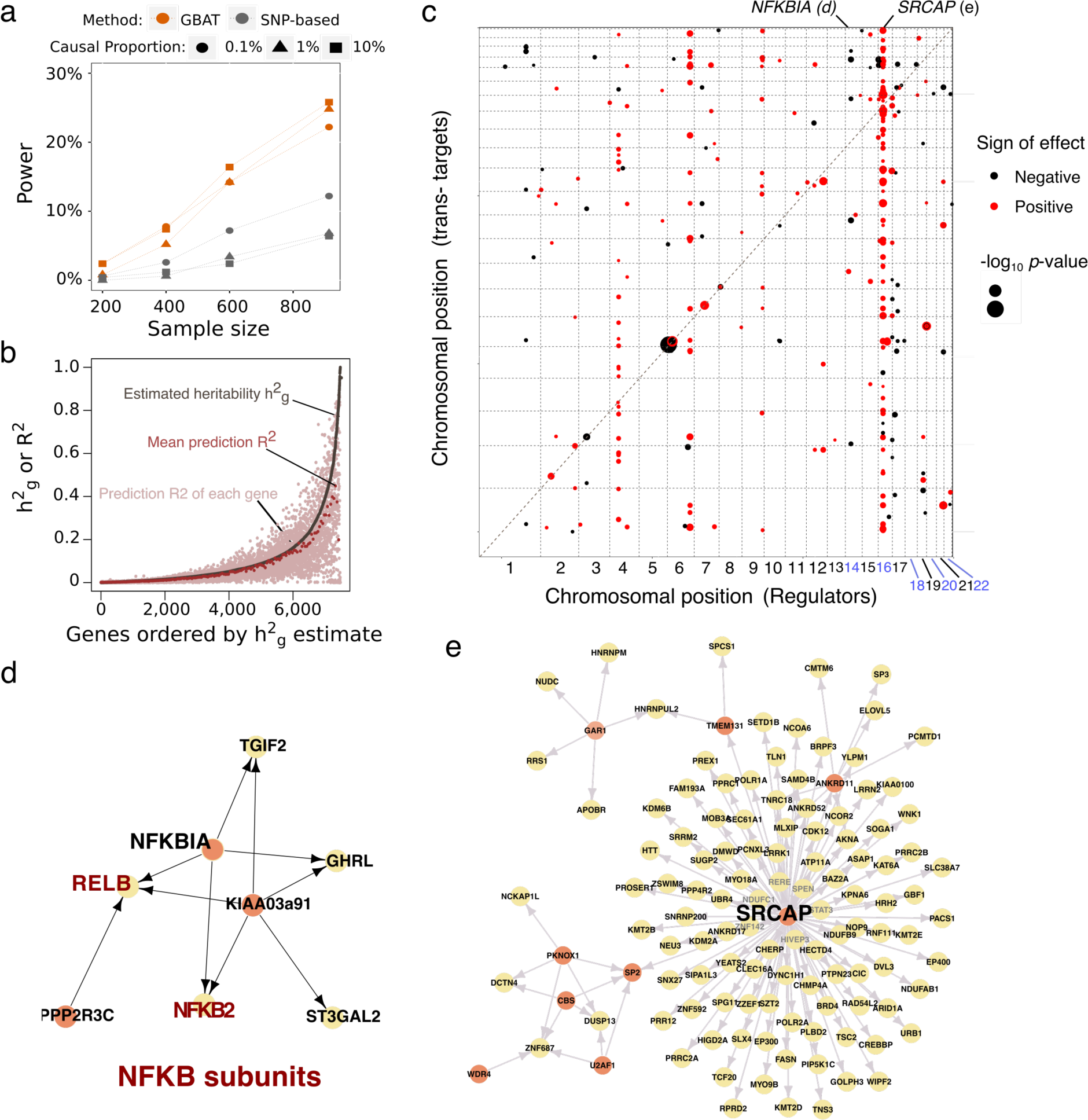
Power of GBAT and *trans-*gene regulation in DGN dataset. a) Power of the GBAT approach in comparison to SNP-based approach. The *cis-*heritability was set to 0.2 and the *trans-*heritability was set to 0.1. Bonferroni correction was used to assess power (accounting for 1 million×15000 SNPs-gene pairs for SNP-based approach and 5,000 (highly heritable gene)×15,000 gene-gene pairs). Power was computed as the fraction of 500 simulations where significant association was identified. Colors represent different methods. b) Comparison of prediction *R*^*2*^ to cis SNP-heritability 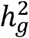. Grey dots denote narrow sense 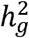 estimated by REML. Red dots denote the mean prediction *R*^*2*^ in bins of 50 genes. Pink dots are the prediction *R*^*2*^ of each gene. c) *Trans* gene regulation signal in DGN at 10% FDR. X-axis are the regulators, and Y-axis are the *trans* target genes whose expression is regulated by the regulators. The size of the dots denote significance of *trans* association (-log10(p-value)). Color of the dots denotes sign of effects. The dashed diagonal line is the x=y line. d)-e) Directed gene network built on significant *trans* signal in DGN for master regulators *NFKBIA* and *SRCAP*. Orange nodes denotes regulators. Yellow nodes are *trans-*genes. Arrows points from regulators to trans-genes.

We next applied GBAT to the DGN dataset to detect *trans-* gene regulation signal for real expression phenotypes. We built cross-validated *cis-*genetic (*CVGP*) expression levels using cvBLUP with variants within 100kb of the transcription start site. Prediction accuracy was assessed using squared correlation (prediction *R*^*2*^) between observed and predicted expression levels. On average across all genes, the prediction *R*^*2*^ is 85% of the *cis* SNP heritability 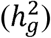 estimated by restricted maximum likelihood (REML) (Figure 2b). We note our prediction accuracy is comparable to prediction methods modeling the sparse genetic architecture of *cis*-gene regulations (Figure 3 of ref^18^ and Figure 4 of ref^19^). After gene-based association test, we computed q-values^20^ from the p-values of all inter-chromosomal gene pairs, and applied the threshold to all inter-and intra-chromosomal gene pairs. At 10% FDR, the final *trans* gene regulation signal consists of 411 regulator-*trans* target gene pairs and 157 unique regulators (Figure 2c, Table S1). Among the 411 *trans* gene pairs, 290 (70.6%) are inter-chromosomal (corresponding to 253 unique inter-chromosomal *trans-*eGenes), and 121(29.4%) are intra-chromosomal (corresponding to 94 unique intra-chromosomal *trans-*eGenes). In contrast, SNP-based eQTL mapping with Matrix eQTL^23^ identified only 90 *trans*-eGenes at 10% FDR in DGN (Supplementary Notes, Table S3). Gene Ontology enrichment analysis by the Database for Annotation, Visualization and Integrated Discovery^21^ (DAVID v6.8, see URLs) showed that the top two enriched categories of the 157 *trans* regulators are DNA binding (Benjamini-Hochberg (BH) FDR = 1.7×10^−4^) and transcription factor activity (BH FDR = 5.3×10^−4^, Table S2). Both regulators and *trans* target genes show heritability enrichment in autoimmune diseases, including lupus, ulcerative colitis, Crohn’s disease and inflammatory bowel disease (Figure S4), by using stratified LDSC^22^.

Among 157 unique *trans* regulators, 20 were found to regulate more than 3 genes (Table S4), supporting the existence of master regulators. For example, we identified *NFKBIA* (NF-Kappa-B Inhibitor Alpha) as a master regulator that regulates expression of four other genes, including two that encode subunits of NF-kappa-B complex: *NFKB2* and *RELB* (Figure 2d). Consistent with the inhibiting effect of NFKBIA on NF-kappa-B subunits, our estimated effect sizes effect size on *NFKB2* and *RELB* are both negative (*NFKB2* beta=-0.19, P=1.0×10^−08^; *RELB* beta=-0.17, P= 2.4×10^−07^). We also identified a master regulator encoded by *SRCAP* on chromosome 16 that regulates 88 other genes (81 are inter-chromosomal signals, Figure 2e). *SRCAP*, short for Snf2 Related CREBBP Activator Protein, encodes the core catalytic component of a chromatin-remodeling complex. *SRCAP* is known to activate the expression of *CREBBP*, consistent with our positive estimated effect in the DGN dataset (effect size=0.25; P= 1.3×10^−14^). The SRCAP complex was shown to regulate key lymphoid fate in haematopoietic system by remodeling chromatin and enhancing promoter accessibility of target genes^23^. In the DGN dataset, the expression level of *SRCAP* is highly correlated with natural killer cell proportion (Pearson correlation = 0.21, P=7.9×10^−11^) and T cell proportions (Pearson correlation=-0.15, P=4.0 x10^−06^). Remarkably, we found that 19 *SRCAP* target genes (out of 88 genes, 2.9 fold enrichment, Fisher’s test) overlap with genes associated with blood cell type proportions (Table S5) from the GWAS catalog (see URLs). Our discovery highlights the gene co-expression network driven by *SRCAP* (Figure 2e) is relevant to haematopoiesis and immune cell type proportions, and suggests that *SRCAP* is a master regulator that controls lineage commitment during haematopoiesis.

GBAT was carefully designed to improve power and reduce false positives in detecting *trans* signals; failing to properly perform the recommended steps led to an increase in false positives or loss of power. For example, incomplete removal of problematic sequence reads resulted in 220% more *trans* signals, most of which are likely to be false positives; correcting for covariates using regression model *E*_*j*_ ∼ *CVGP*_*i*_ + *SV*_*i*_ resulted in 39% more *trans* signals, also likely due to false positives inflation (Supplementary notes).

GBAT can be used to detect other *trans* gene regulatory events, such as splicing, methylation, protein regulation or gene network response to stimulus. As larger studies become available for increasingly diverse populations, tissues, and functional genomic measurements, we foresee more *trans* regulation discoveries will reveal new disease genes and mechanisms.

## Methods

### Cross-validated *cis*-genetic prediction with cvBLUP

The cross-validated prediction by cvBLUP is a cross validated version of a standard linear mixed model (LMM) prediction, or best linear unbiased predictor (BLUP). We consider an LMM as below:

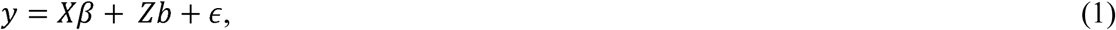

where y is the phenotype, in particular the expression of gene, measured on N individuals. X is a matrix of covariates, including an intercept. Z is a standardized *N*×*M* matrix of M SNPs within the *cis* region of the gene. *b* is the vector of effect sizes for the SNPs in *Z*, modeled as normally distributed by 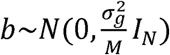. The total *cis*-genetic contribution to the phenotype is then the product *Zb*, with distribution 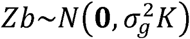, where *K* is the genetic relationship matrix defined as 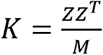. Finally, *ϵ* is a vector of non-genetic effects, modeled as 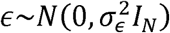. Phenotype y therefore has the distribution: *y*∼*N*(*Xβ, V*), with 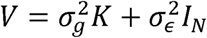. We use standard REML to get estimates of the LMM variance components, 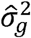 and 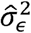. The estimate of the narrow sense heritability 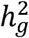 is then the ratio of estimated genetic variance to total variance: 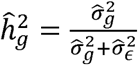.

The BLUPs for the random effects are: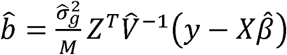, and the genetic predictor, or fitted-value of y is calculated as 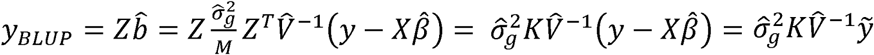, where 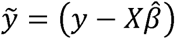 are the phenotypic residuals after removing the contributions of the covariates X.

The standard BLUPs, *yBLUP*, over-fit the training data, meaning that they are highly correlated with the noise term *ϵ*. Cross-validation is often used to mitigate overfitting. In our analysis, we use a leave-one-out cross-validation scheme to generate a set of out-of-sample LMM predictions: each subject is left out of the dataset in turn; the remaining subjects are used to estimates 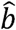, and then the genetic contribution to the left-out subject’s phenotype is defined using 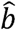.

The resulting collection of cross-validated LMM estimates, or cvBLUPs, is still a strong estimator of the true *cis*-genetic contribution to the phenotype, but does not have spurious correlations with *ϵ*. Fortunately, leave-one-out cross-validation is mathematically simple for BLUPs. Given 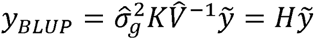, such that the prediction is a linear operator 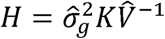 applied to 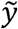, the out-of-sample prediction and prediction errors can be simply calculated from a single model fit. The cvBLUPs can therefore be calculated in linear time as:

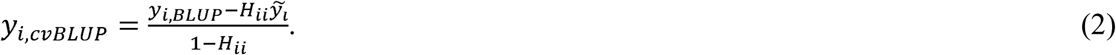

For any gene *i*, the cross validated *cis*-genetic prediction value (*CVGP*_*i*_) is calculated with all *cis* genetic variants (±100kb to transcription starting sites) of gene *i* by using Equation (2).

### Gene-based association testing of *trans* signals (GBAT)

For any gene *i*, we test the gene-based *trans* association between *CVGP*_*i*_ and the expression level of gene j (*E*_*j*_). Gene *j* is in *trans* to gene *i* if it is at least 1Mb away from gene *i*. To improve association power and reduce spurious *trans* association signal, we used supervised SVA conditioning on *CVGP*_*i*_ (*SV*_*i*_) as covariates^11, 14, 24^. We first regressed out *SV*_*i*_ from quantile normalized expression *E*_*j*_ in a linear model *E*_*j*_ *∼ SV*_*i*_. Then we used the quantile normalized residuals *E j* ^*’*^ to test for trans association: *E j* ^*’*^*∼ CVGP_i_.*

To compute the FDR levels from all *trans* association tests, we used only the summary association p-values from the inter-chromosomal trans association tests. We used 10% FDR for significant *trans* signals. We further removed gene pairs that are cross mappable due to sequence similarities surrounding the gene pairs^25^.

## URLs

GBAT: the pipeline will be available before publication

PLINK 2.0: https://www.cog-genomics.org/plink/2.0/

Michigan Imputation Server: https://imputationserver.sph.umich.edu/index.html

ENCODE 36 k-mer of the reference human genome: http://hgdownload.cse.ucsc.edu/goldenPath/hg19/encodeDCC/wgEncodeMapability/wgEncodeCrgMapabilityAlign36mer.bigWig

DAVID 6.8: https://david.ncifcrf.gov/tools.jsp

Matrix eQTL: http://www.bios.unc.edu/research/genomic_software/Matrix_eQTL/

GWAS Catalog: https://www.ebi.ac.uk/gwas/

## Acknowledgements

We thank members of Zaitlen lab, Price lab and Y Li for helpful discussions. This work was supported by R01 MH115676.

## References

1. Franzén, O. et al. Cardiometabolic risk loci share downstream cis- and trans-gene regulation across tissues and diseases. Science 353, 827–830 (2016).

2. Rau, C. D. et al. Systems Genetics Approach Identifies Gene Pathways and Adamts2 as Drivers of Isoproterenol-Induced Cardiac Hypertrophy and Cardiomyopathy in Mice. Cell Syst 4, 121–128.e4 (2017).

3. Small, K. S. et al. Regulatory variants at KLF14 influence type 2 diabetes risk via a female-specific effect on adipocyte size and body composition. Nat Genet 50, 572–580 (2018).

4. Sun, B. B. et al. Genomic atlas of the human plasma proteome. Nature 558, 73–79 (2018).

5. Price, A. L. et al. Single-tissue and cross-tissue heritability of gene expression via identity-by-descent in related or unrelated individuals. PLoS Genet 7, e1001317–9 (2011).

6. Grundberg, E. et al. Mapping cis- and transregulatory effects across multiple tissues in twins. Nat Genet 44, 1084–1089 (2012).

7. Liu, X. et al. Functional Architectures of Local and Distal Regulation of Gene Expression in Multiple Human Tissues. The American Journal of Human Genetics 100, 605–616 (2017).

8. Westra, H.-J. et al. Systematic identification of trans eQTLs as putative drivers of known disease associations. Nat Genet 45, 1238–1243 (2013).

9. Battle, A. et al. Characterizing the genetic basis of transcriptome diversity through RNA-sequencing of 922 individuals. Genome Research 24, 14–24 (2014).

10. Stegle, O., Parts, L., Piipari, M., Winn, J. & Durbin, R. Using probabilistic estimation of expression residuals (PEER) to obtain increased power and interpretability of gene expression analyses. Nature Protocols 2012 7:3 7, 500–507 (2012).

11. Leek, J. T. & Storey, J. D. Capturing Heterogeneity in Gene Expression Studies by Surrogate Variable Analysis. PLoS Genet 3, e161–1735 (2007).

12. Kang, H. M., Ye, C. & Eskin, E. Accurate discovery of expression quantitative trait loci under confounding from spurious and genuine regulatory hotspots. Genetics 180, 1909–1925 (2008).

13. Joo, J. W. J., Sul, J. H., Han, B., Ye, C. & Eskin, E. Effectively identifying regulatory hotspots while capturing expression heterogeneity in gene expression studies. Genome Biol 15, r61 (2014).

14. Dahl, A., Guillemot, V., Mefford, J., Aschard, H. & Zaitlen, N. Adjusting For Principal Components Of Molecular Phenotypes Induces Replicating False Positives. (2017). doi:10.1101/120899

15. GTEx Consortium. Genetic effects on gene expression across human tissues. Nature 550, 204–213 (2017).

16. Zhou, X., Carbonetto, P. & Stephens, M. Polygenic Modeling with Bayesian Sparse Linear Mixed Models. PLoS Genet 9, e1003264 (2013).

17. Zou, H. & Hastie, T. Regularization and variable selection via the elastic net. Journal of the Royal Statistical Society: Series B (Statistical Methodology) 67, 301–320 (2005).

18. Gamazon, E. R. et al. A gene-based association method for mapping traits using reference transcriptome data. Nat Genet 47, 1091–1098 (2015).

19. Gusev, A. et al. Integrative approaches for largescale transcriptomewide association studies. Nat Genet 48, 245–252 (2016).

20. Storey, J. D. & Tibshirani, R. Statistical significance for genomewide studies. PNAS 100, 9440–9445 (2003).

21. Huang, D. W., Sherman, B. T. & Lempicki, R. A. Bioinformatics enrichment tools: paths toward the comprehensive functional analysis of large gene lists. Nucl. Acids Res. 37, 1–13 (2009).

22. Finucane, H. K. et al. Partitioning heritability by functional annotation using genome-wide association summary statistics. Nat Genet 47, 1228–1235 (2015).

23. Ye, B. et al. Suppression of SRCAP chromatin remodelling complex and restriction of lymphoid lineage commitment by Pcid2. Nat Commun 8, 1518 (2017).

24. Chen, J. et al. Fast and robust adjustment of cell mixtures in epigenome-wide association studies with SmartSVA. BMC Genomics 18, 413 (2017).

25. Saha, A. et al. Co-expression networks reveal the tissue-specific regulation of transcription and splicing. Genome Research 27, 1843–1858 (2017).

